# Multiscale Biomechanical and Electrophysiological Modeling of Nociceptive Activation in Musculoskeletal Joint Disorders: Insights from the Temporomandibular Joint

**DOI:** 10.64898/2026.07.17.738969

**Authors:** Jian Chen, Shuchun Sun, Farhad Ahmadi, Peng Chen, Jiaxin Chai, Jichao Zhao, Brooke Damon, Konstantinia Almpani, Janice Lee, Hai Yao

## Abstract

**Background and objective:** Musculoskeletal joint disorders often show inconsistent relationships between structural degeneration and nociceptive pain. Temporomandibular joint (TMJ) disc displacement represents a clinically relevant model for investigating the structure-function-pain relationship. This study aimed to develop a multiscale computational framework integrating biomechanics, three-dimensional (3D) neural morphology, and electrophysiology to quantitatively link TMJ structural alterations, biomechanical loading, and peripheral nociceptive activation.

**Methods:** Strain distributions in the TMJ disc and retrodiscal tissue during mouth opening and clenching were computed in ArtiSynth under varying degrees of displacement. Human TMJ 3D nerve architecture was reconstructed using porcine TMJ nerve mapping data as an anatomical reference, and ion channel dynamics were implemented in NEURON. Model coupling was achieved by mapping biomechanical strain fields onto nociceptor membranes to simulate mechanosensitive currents and action potential propagation to the trigeminal ganglion.

**Results:** Anterior DDwoR induced a severity dependent strain pattern in the TMJ disc and retrodiscal tissue, including posterior redistribution, increased strain magnitude, prolonged activation, and broader retrodiscal tissue involvement. Displacements of 4, 6, and 8 mm produced larger mechanosensitive currents, broader terminal depolarization, and higher trigeminal firing rates during mouth opening (6, 18, and 28 Hz) and clenching (8, 20, and 28 Hz), whereas 0- and 2-mm displacements produced negligible neural activation.

**Conclusions:** This study establishes a multiscale biomechanical-electrophysiological framework linking TMJ structural alterations to peripheral nociceptive activation. The framework quantitatively connects macroscale strain patterns with microscale neural activation, suggesting that anterior disc displacement may amplify peripheral nociceptive signaling by increasing the overlap between elevated strain and densely innervated retrodiscal tissue.

## Introduction

Musculoskeletal joint disorders impair physical function and quality of life and are a major source of pain and disability [1, 2]. However, the relationship between structural degeneration and pain remains poorly defined, as pain may occur in the absence of detectable structural alterations, while pronounced structural degeneration may exist without pain [3]. Consequently, structural alterations observed on computed tomography (CT) or magnetic resonance imaging (MRI) scans are inadequate for the explanation of the pain symptomatology [4]. The temporomandibular joint (TMJ), a load-bearing synovial joint subjected to repetitive mechanical loading during different oral functions such as mastication and speech, represents a clinically relevant example of this structure-function-pain discordance [3, 5]. Temporomandibular disorders (TMDs) encompass a spectrum of conditions affecting the TMJ pain, associated masticatory muscles, and surrounding jaw tissues, impacting more than 11 million adults in the U.S. [6, 7]. Among various TMJ disorders, anterior TMJ disc displacement is one of the most prevalent and clinically impactful subtypes [8], particularly disc displacement without reduction (DDwoR), which accounts for approximately 36% of patients [9]. This structural alteration, in which the articular disc is displaced and fails to return to its normal position during mandibular movement, gives rise to pain symptoms that constitute the primary reason for seeking medical care [10, 11].

Pain involves the interpretation and perception of nociceptive signals triggered by noxious mechanical, thermal, or chemical stimuli acting on the terminals of pain-sensing nerve fibers (nociceptors), and its intensity generally corresponds to the magnitude of the stimulus [12]. In TMJ DDwoR, structural alterations lead to abnormal joint biomechanics, resulting in aberrant mechanical stimuli acting on nociceptors and influencing nociceptive signal generation. This process involves multiscale and multiphysics interactions, as illustrated in **Fig. 1**. Determining causal relationships among these factors remains challenging in in vivo models. Computational modeling as an effective tool can quantitatively analyze the relationships linking human TMJ structural alterations to biomechanical loading [13, 14] and subsequent nociceptive signaling. Nevertheless, current TMJ modeling studies have focused separately on macroscopic joint biomechanics and microscopic neural mechanisms, thereby lacking an integrated approach. Therefore, a multiscale computational model integrating TMJ structure, biomechanics, and nociceptive signal activation is needed to investigate how altered joint biomechanics may contribute to peripheral nociceptive activation in TMDs.

**Fig. 1.**
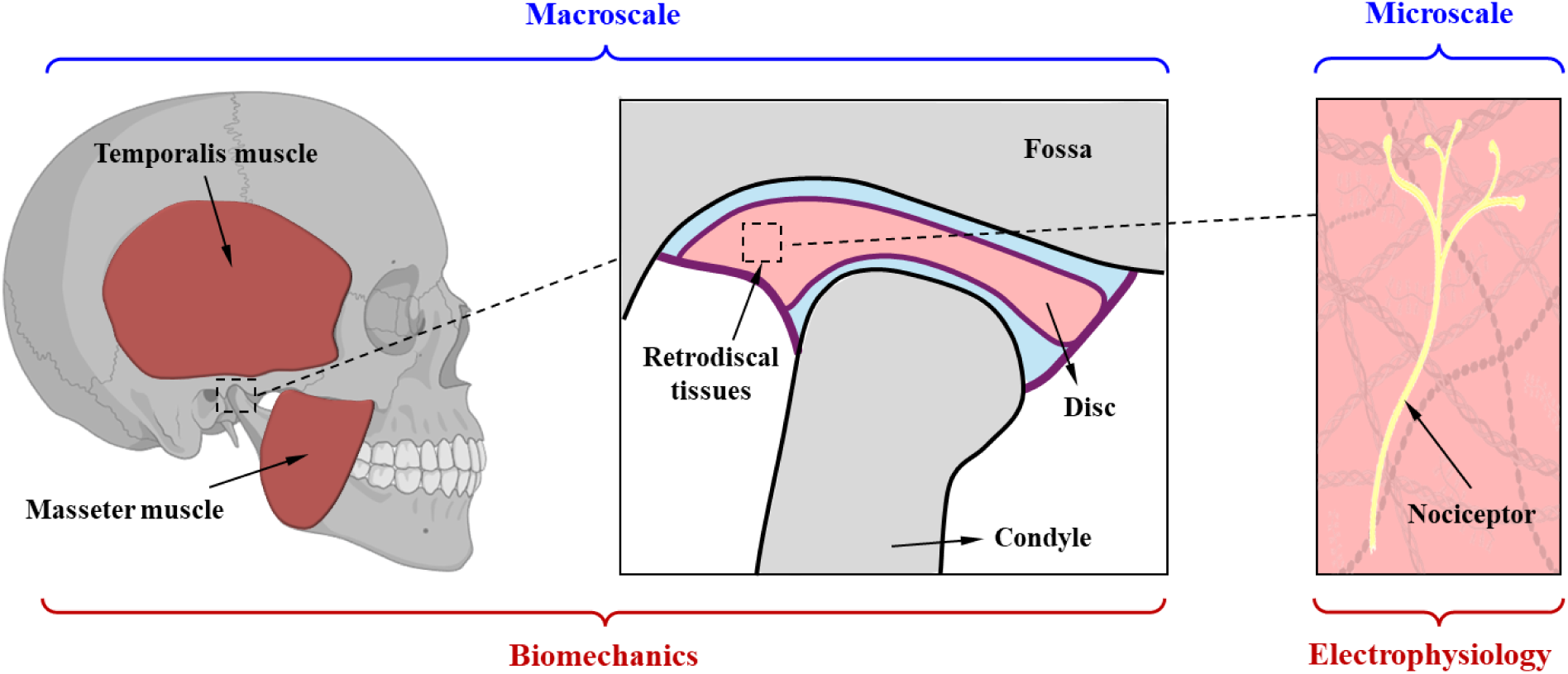
Multiscale and multiphysics characteristics of the human TMJ system: At the macroscale, the system consists of the musculoskeletal components of the jaw, including the skull, mandible, muscles, TMJ disc, and retrodiscal tissue, which govern joint motion and tissue deformation. Embedded within these soft tissues, the microscale component consists of nociceptive nerve fibers and terminals, which convert local mechanical strain into electrophysiological signals.

At the macroscale, TMJ biomechanical models based on finite element method integrate musculoskeletal geometry, material properties, and loading conditions to estimate mechanical stimuli, including stress, strain, and contact forces, that may drive nociceptor activation at the microscale [15–17]. Under TMJ disc displacement conditions, biomechanical models have revealed alterations in joint biomechanics across functional tasks. During jaw opening, anterior disc displacement altered disc stress distribution, increased connective tissue stress, and elevated the articular surface friction [18], with the region of maximum disc stress shifting from the intermediate zone to the posterior band [19]. During maximum clenching, anterior disc displacement altered TMJ stress distribution and increased shear stresses in the articular disc and bilaminar zone [20]. These findings indicate that DDwoR modifies joint biomechanics, particularly affecting the disc and retrodiscal tissue. However, although biomechanical analyses often suggest a potential link between altered joint structure and pain, they do not quantitatively link structural changes, local mechanical loading, nerve distribution, and nociceptor activation, leaving the mechanistic pathway from DDwoR to nociceptive pain unresolved.

To address this gap, the macroscale biomechanical outputs should be integrated with microscale electrophysiological modeling of nociceptor mechanotransduction, which converts mechanical loading into electrical signals. Establishing such a model requires both the spatial distribution and electrophysiological properties of TMJ innervation. TMJ innervation arises primarily from the trigeminal ganglion [21, 22], and previous histological and imaging studies indicate that the human TMJ disc and its surrounding tissues exhibit region-specific innervation: the central region is largely devoid of nerves, whereas nerves are predominantly concentrated in the peripheral regions, with greater nerve density in the anterior and posterior regions [23]. However, most existing evidence is based on two-dimensional (2D) histological sections, which provide limited spatial information and cannot fully capture three-dimensional (3D) neural architecture or neural connectivity across TMJ tissues. Electrophysiological studies further indicate that unmyelinated C-fibers constitute the predominant nociceptive fiber type innervating the retrodiscal tissues in both healthy individuals and TMD patients [24, 25]. Patch-clamp recordings have demonstrated that mechanosensitive ion channels in peripheral nociceptors play a critical role in converting mechanical deformation into nociceptive activity [26–29]. Computational models incorporating ion-channel dynamics have been successfully used to investigate nociceptive signaling, in cutaneous [30, 31], visceral [32, 33], lower limbs [34] and dental area [35], demonstrating the utility of biophysical modeling for linking mechanical, thermal and electrical stimuli to neural activation. However, these models generally focus on single-fiber activation and do not incorporate the spatial organization of neural networks within TMJ tissues. Therefore, the lack of an integrated 3D neural architecture and electrophysiological model limits the mechanistic understanding of how TMJ biomechanics contributes to nociceptor activation.

The objective of this study was to develop a multiscale computational framework using the TMJ as a representative model system to quantitatively link joint structural alterations with peripheral nociceptive activation. The framework integrates macroscale TMJ biomechanical modeling with microscale 3D nociceptor morphology and electrophysiology to predict how strain patterns generated during mouth opening and clenching are converted into action potential firing at the trigeminal ganglion. Using this framework, we investigated how different degrees of anterior DDwoR alter regional strain distributions and drive nociceptive signaling. Although developed for the TMJ, this framework may provide a basis for future investigations of structure-function-pain relationships in other musculoskeletal joint disorders.

## Materials and methods

### 2.1 Biomechanical Model

To investigate the mechanical stimuli applied to the TMJ disc and retrodiscal tissue during mouth opening and clenching under varying degrees of TMJ anterior DDwoR, we developed integrated rigid body dynamics and finite element models in the ArtiSynth Modeling Toolkit [36], as illustrated in **Fig. 2A**.

**Fig. 2.**
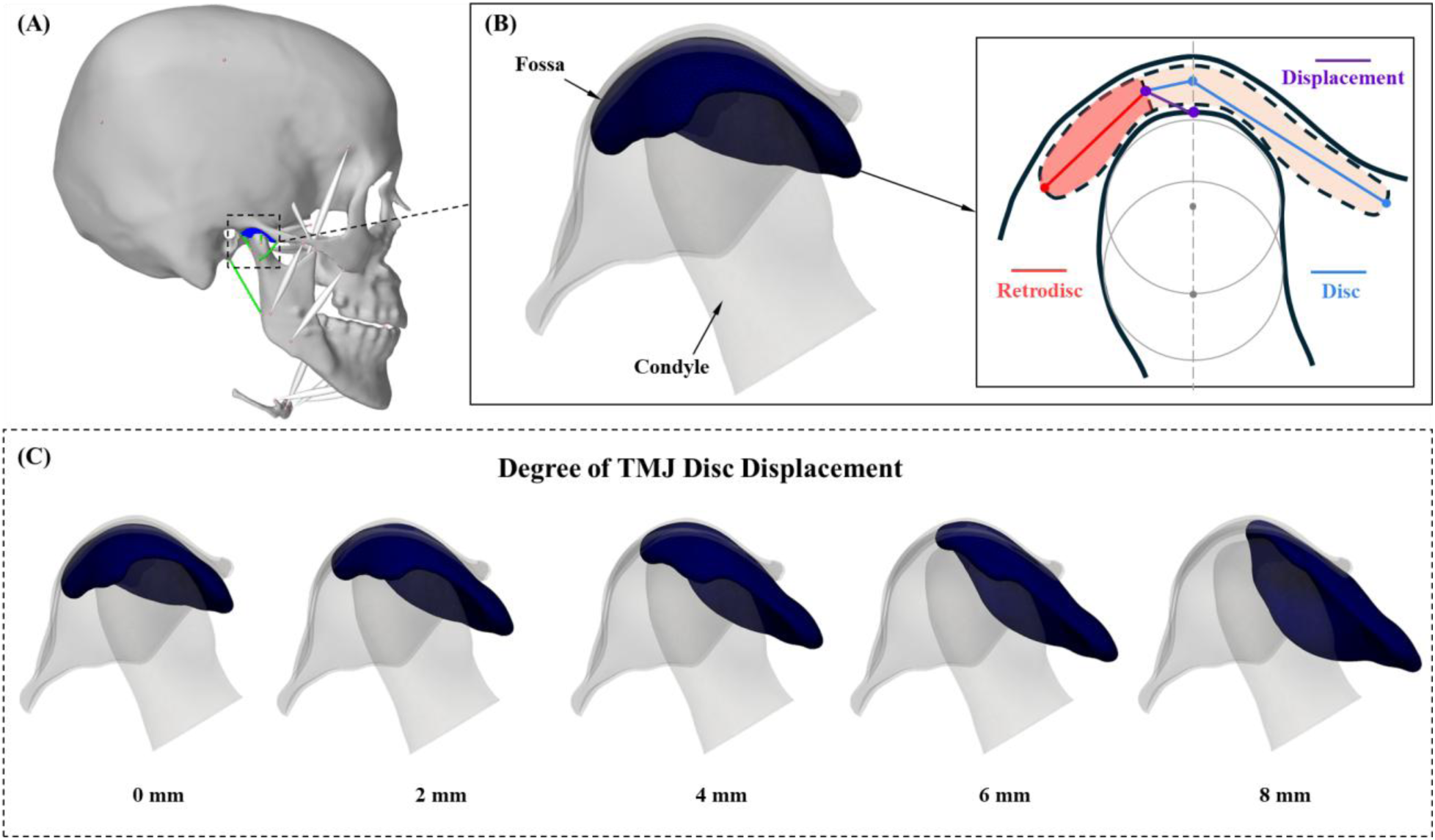
Schematic representation of TMJ anterior disc displacement without reduction (DDwoR): (A) Musculoskeletal model of the human jaw implemented in ArtiSynth; (B) The definition of disc displacement degree relative to the condyle; and (C) TMJ models with varying severities of anterior disc displacement in the right joint.

The skull, mandible and hyoid bone geometries were reconstructed from full head cone-beam computed tomography (CBCT) scans, and the TMJ disc and retrodiscal tissue were segmented from head MRI scans to construct a generic healthy TMJ model. The study adhered to ethical standards and was approved by the Institutional Review Board. All the bones were modeled as rigid bodies, with the skull fixed and the mandible allowed to move. For clenching, the hyoid bone was fixed throughout the simulation. For mouth opening, the hyoid bone was pre-positioned 7 mm posteriorly and 7 mm inferiorly before task initiation and then fixed throughout the simulation, approximating physiological hyoid displacement during maximum mouth opening [37, 38]. The TMJ disc and retrodiscal tissue were defined as finite element bodies and modeled as the Mooney-Rivlin hyperelastic materials. Articular cartilages were simulated as elastic foundations. Four cable constraint groups were used to represent simplified disc attachments in the anterior, medial, lateral, and posterior directions. The anterior, medial, and lateral attachments connected the disc to the condyle, whereas the posterior attachment connected the skull to the disc. Three anatomical TMJ ligaments were also included: the lateral temporomandibular, sphenomandibular, and stylomandibular ligaments. Twelve Hill-type point-to-point muscles were implemented, including the posterior, medial, and anterior parts of the temporalis muscle; the superior and inferior heads of the lateral pterygoid muscle; the superficial and deep masseter muscles; the medial pterygoid muscle; the anterior digastric muscle; the geniohyoid muscle; and the anterior and posterior mylohyoid muscles. A schematic overview of all components included in the musculoskeletal finite element model is provided in **Supplementary Figure 1**. Detailed parameters were summarized in the **Table 1** and **Supplementary Note 1**.

**Table 1.** Biomechanical Model Components and Parameters.

| Components | Constitutive model | Description |
| --- | --- | --- |
| TMJ disc, retrodiscal tissue [38, 45] | Mooney-Rivlin strain energy density:<br>$W(I_1, I_2) = C_1(I_1 - 3) + C_2(I_2 - 3)$ | $C_1=0.9$ MPa, $C_2=0.0009$ MPa for the disc; $C_1=0.27$ MPa, $C_2=0.00027$ MPa for the retrodiscal tissue. |
| Articular cartilages [38, 45] | Elastic foundation model describing the contact pressure $p$ between the articular cartilage and the TMJ disc as a function of penetration depth $d$:<br>$p(d) = K \ln \left( 1 - \frac{d}{h} \right), K \equiv \frac{-(1-\nu)E}{(1+\nu)(1-2\nu)}$ | With cartilage thickness $h=0.4$ mm, elastic modulus $E=2.7$ MPa, and Poisson's ratio $\nu=0.49$. |
| Disc attachments and ligaments [38, 45] | Elastic cable behavior with slack lengths of 4 mm, 1.9 mm, 1.9 mm, 7.5 mm for the anterior, medial, lateral and posterior disc attachments, respectively. Slack lengths of 4 mm, 0 mm, 0 mm for lateral temporomandibular, sphenomandibular, and stylomandibular ligaments, respectively | The elongation stiffness of 250 MPa applied once the tendon slack length was exceeded |
| Muscles [38, 45] | Hill-type muscle: the total force generated by each muscle is calculated as the sum of active contractile force and passive elastic force | The detailed parameters were summarized in the **Supplementary Note 1**. |

Different degrees of disc displacement were defined based on established criteria [39, 40]. As shown in **Fig. 2B**, the long axis of the condyle was determined using a two-step method, and disc displacement relative to the condyle was measured as the distance between the intersection of this axis with the condylar bone surface and the most posterior point of the disc. According to MRI observations of anterior DDwoR [41, 42], disc displacement distances corresponding to the healthy condition (0-mm displacement) and anterior disc displacement states (2-, 4-, 6-, and 8-mm displacements) were defined in **Fig. 2C**. For all displacement cases, the skull and mandible were fixed in the same reference closed-mouth position. A prescribed displacement was applied to the anterior disc attachment region to pull the deformable finite element disc to the target anterior displacement distance. During this process, disc-condyle and disc-fossa contact interactions were maintained, allowing the disc to deform and conform to the surrounding articular surfaces. The resulting disc geometry was exported as the initial configuration for subsequent functional simulations. For the mouth opening and clenching tasks, muscle activation levels were summarized in the **Supplementary Figure 2**. In both tasks, the target activation level was reached at 0.25 s and then maintained until the end of the simulation at 0.5 s.

The model was validated by comparing the simulated mandibular motion with experimentally acquired kinematic data recorded under healthy conditions without disc displacement using custom tracking devices [43], and by comparing simulated mandibular motion across varying degrees of disc displacement. The validation included assessments of mandibular position and configuration during maximum mouth opening, central incisor displacement, and rotation magnitude. Quantitative analysis showed that the model captured the endpoint opening configuration and overall temporal trends of mandibular motion, with small endpoint errors for central incisor displacement and rotation magnitude and strong curve correlations, although moderate discrepancies remained during the intermediate phase of mouth opening. Mandibular motion remained nearly unchanged across different severities of disc displacement. Details of the validation procedures and quantitative analysis are summarized in **Supplementary Materials Note 2**.

Considering the large deformations [44], the spatial patterns of equivalent logarithmic strain were evaluated using the following formulation:

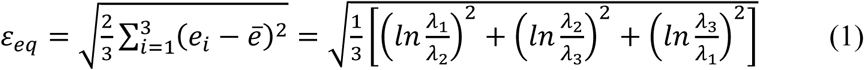

where *λ*_i_ (*i* = 1, 2, 3) are the principal stretch ratios, *e*_i_ = *ln*(*λ*_i_), 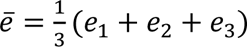. This strain value will be used to map onto the membranes of nociceptors at the microscale.

### 2.2 Coupling Between the Biomechanical and Electrophysiological Models

Before coupling the biomechanical model with the electrophysiological model, we constructed a microscopic nociceptive nerve network distributed within the macroscopic TMJ disc and retrodiscal tissue. Nerve location, density, and morphological features were derived from tissue clearing and 3D nerve mapping of porcine specimens [46], with image segmentation of representative subdomains revealing bifurcating terminal fiber structures (**Fig. 3A**). These data informed the reconstruction of human TMJ innervation geometry, including nodal coordinates, connectivity, and diameters. Soma representing the trigeminal ganglion were positioned superior to the human TMJ disc according to the innervation patterns [21, 22]. Peripheral axons and terminal branches were generated using a randomized Rapidly Exploring Random Tree (RRT) algorithm [47] constrained to the innervated TMJ regions, including the disc periphery and throughout the retrodiscal tissue. Probabilistic branching was introduced to emulate dendritic arborization [48], while constraints on total fiber length [46] and full connectivity to the soma ensured physiologically plausible nerve trajectories. The reconstructed nerve network was represented as an independent topology of connected nodes and segments geometrically embedded within the tissue domain, and did not contribute to the finite element mechanical response and was discretized separately from the tissue mesh, as shown in **Fig. 3B**.

**Fig. 3.**
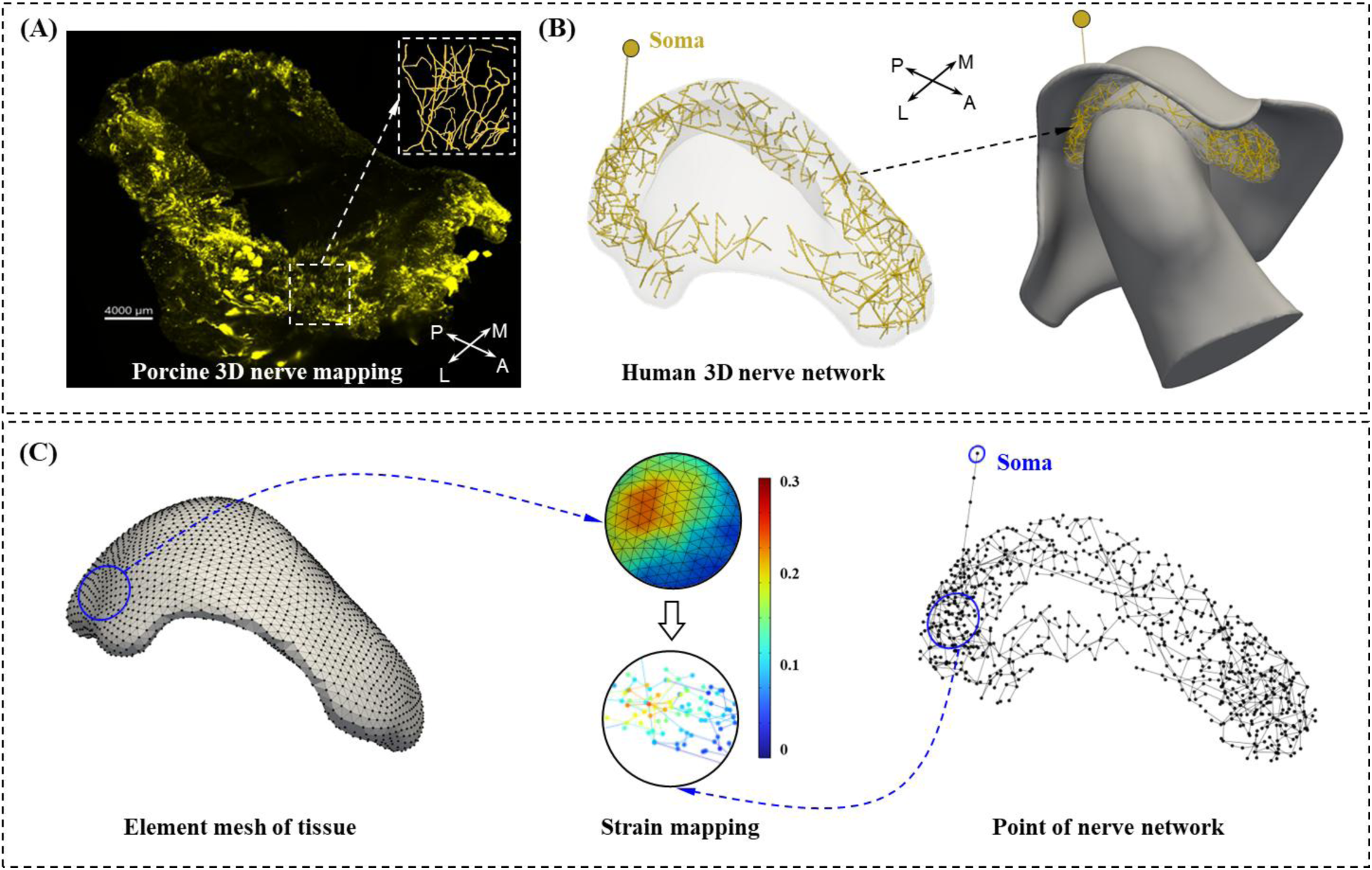
Reconstruction of the nociceptive nerve network and multiscale mapping of tissue strain onto embedded nociceptors: (A) 3D innervation imaging of porcine TMJ disc and retrodiscal tissue; (B) Reconstructed microscopic nociceptive nerve network within the human TMJ disc and retrodiscal tissue, including trigeminal ganglion soma, peripheral axons, and terminal branches; and (C) Multiscale coupling strategy in which equivalent strain fields from the macroscopic finite element tissue mesh were spatially interpolated onto microscopic nerve network nodes and used as nociceptive mechanical inputs for the electrophysiological simulations.

After the nerve architecture was established, the equivalent strain field from the macroscopic tissue mesh was used as the source of nociceptive mechanical stimulation for the microscopic nerve network, as shown in **Fig. 3C**. Strain at each nerve node was obtained by spatial interpolation of the surrounding tissue mesh node strains, and the resulting time-varying strain inputs were sequentially applied during the following electrophysiological simulations. The interpolated local strain modulated the open probability of mechanosensitive channels and thereby generated *I*_MS_, the ionic current driving neural activation in the subsequent electrophysiological model [29, 32, 49, 50]:

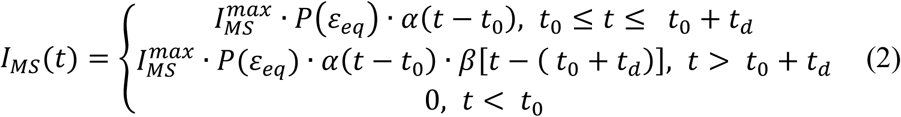

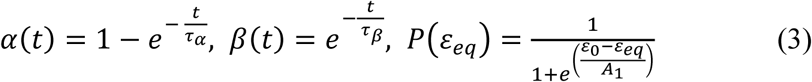

where *t*_0_ and *t_d_* denote the onset time and duration of the stimulus, respectively. 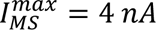 is the maximum ionic current of the mechanosensitive channel, *α*(*t*) and *β*(*t*) are the activation and inactivation functions with *τ_α_* = 50 *ms* and *τ_β_* = 400 *ms* as the activation and inactivation time constants, respectively. *P*(*ε_eq_*) represents the strain-dependent open probability of the mechanosensitive channel, modeled as a Boltzmann-type gating function, *ε_eq_* is the strain on membrane, *A*_1_ = 0.02 is the gating parameters, and *ε*_0_ = 0.1 represents the half-activation strain threshold.

### 2.3 Neural electrophysiological Model

After reconstructing the nerve fiber geometry and coupling it with tissue derived mechanical stimuli, ion channel dynamics were incorporated to simulate neural responses. The nociceptor membrane was modeled using Hodgkin-Huxley type ion channel dynamics:

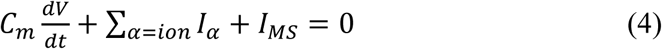

where *V* is the nociceptor membrane potential, *C_m_* is the membrane capacitance per unit area, *I_α_* denotes the ionic current driven by electrochemical potential gradients and governed by channel conductance, and *I_MS_* is the ionic current of the strain-activated mechanosensitive ion channels.

As shown in **Fig. 4**, each nerve fiber was divided into three functional compartments: a transducer zone, a spike initiation zone, and a passive zone [51]. In the transducer zone, mechanosensitive ion channels were gated by local membrane strain derived from finite element simulations, generating ionic currents that induced local depolarization and propagated electrically to the spike initiation zone. There, active ion channels converted the depolarization into action potential. The density of active ion channels gradually increased from the transducer zone to the spike initiation zone, while the passive zone supported action potential propagation and axial ionic diffusion. Segment lengths were assigned according to compartment function: 2.5 μm in the transducer zone, 1 μm in the spike initiation zone, and 5 μm in the passive zone [50].

**Fig. 4.**
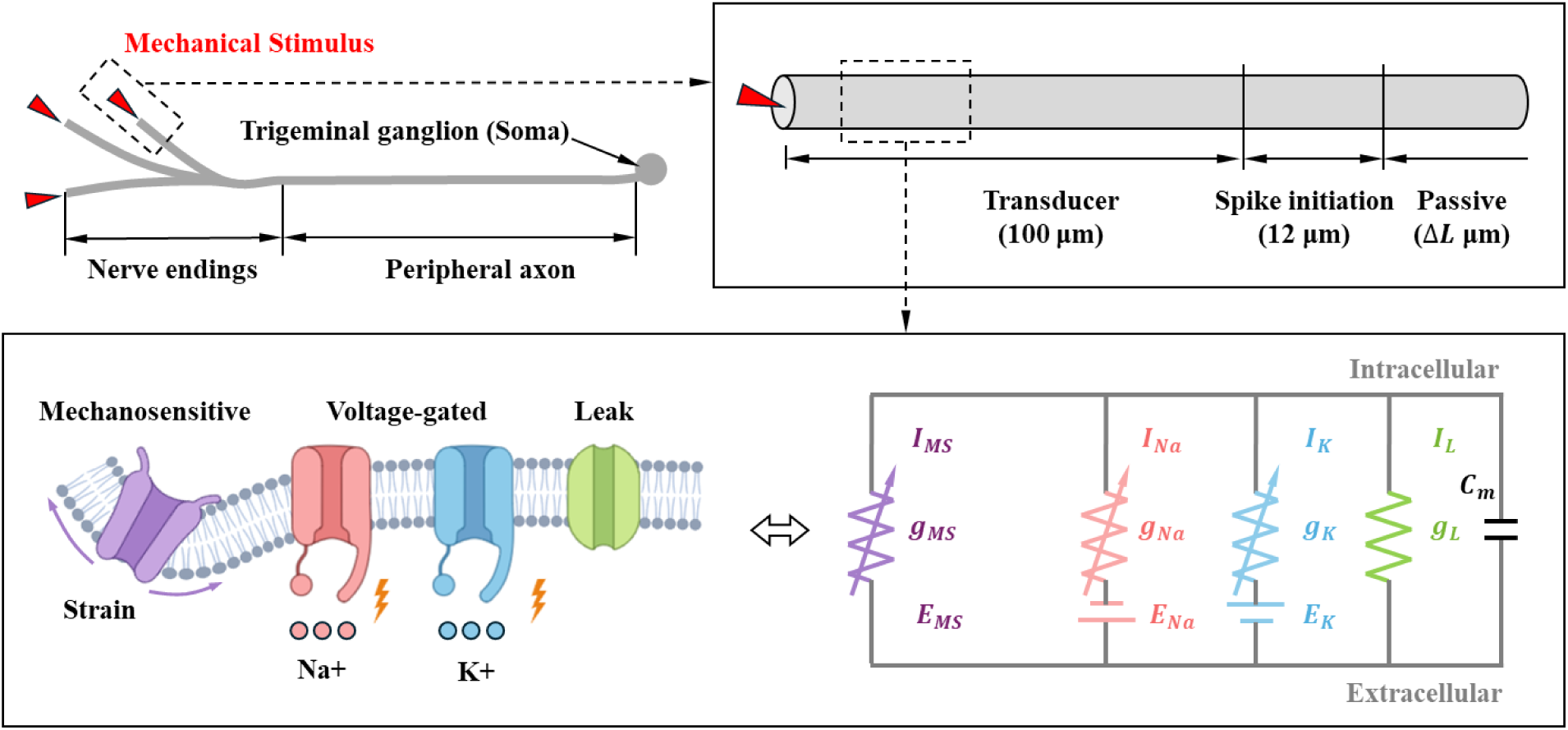
Electrophysiological membrane model of the nociceptor: Peripheral nerve endings were divided into transducer, spike initiation, and passive zones. In the transducer zone, the membrane model incorporated strain-activated mechanosensitive, voltage-gated sodium and potassium, and leak ion channels represented by an equivalent electrical circuit.

For the passive membrane properties, the passive conductance equilibrium potential was set to −60 mV, yielding a simulated resting potential of −60 mV [52]. Membrane capacitance of 1 *μFcm*^−2^ was set for all compartments. Passive membrane resistance of 10000 *Ωcm*^−2^ was set for all compartments in the passive zone [53]. The axial resistance *R_a_* in all compartments, apart from the transducer zone, was 150 *Ωcm* [50]. The *R_a_* of transducer zone was set to 2.25 *MΩcm* which has been demonstrated that the insertion of mitochondria into the dendrite increases *R_a_* of the dendrite 15-fold [54].

For active membrane properties, the nociceptor model incorporated Hodgkin-Huxley-type ion channels [50], including four sodium channel types: Tetrodotoxin (TTX)-sensitive sodium channels, TTX-sensitive persistent sodium channels, Nav1.9 TTX-resistant sodium channels, and Nav1.8 TTX-resistant sodium channels [55, 56]; Three types of potassium channels included: delayed rectifier potassium channels, A-type potassium channels, Kv7/M channels [56–59]; One type of mixed cation channel: hyperpolarization-activated channels [59]. Detailed parameters of ion channels were provided in the **Supplementary Materials Note 3**. Action potential simulations were performed in the NEURON simulation environment [60] with a fixed time step of 0.025 ms over a total simulation duration of 550 ms. Under the one-way sequential coupling strategy, the electrophysiological response does not feed back into the biomechanical simulation.

## Results

### 3.1 Strain Distributions in TMJ Disc and Retrodiscal Tissue

The relative positional changes among the fossa, disc, and condyle at 0.25s during maximum mouth opening and clenching are shown in **Figs. 5A and 6A**, respectively. The corresponding equivalent strain distributions are presented in **Figs. 5B and 6B**. The healthy condition showed the most centralized strain distribution. As anterior DDwoR severity increased, elevated strain progressively shifted posteriorly during both tasks. Disc morphology also varied with displacement severity, as the disc conformed to the condylar head during loading. Severe anterior DDwoR (≥4 mm) produced visibly flatter disc shapes than lower displacement levels.

**Fig. 5.**
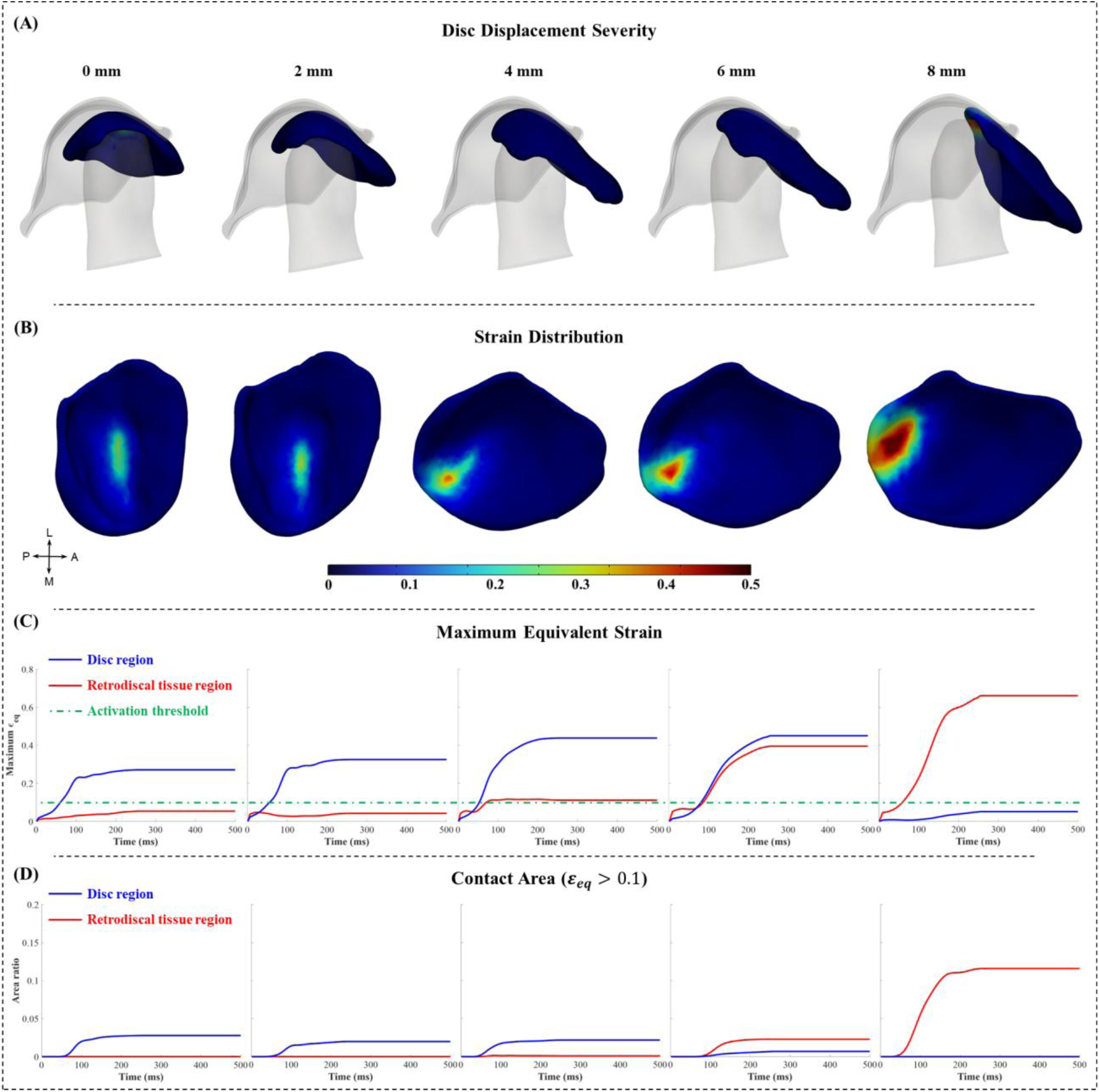
Strain distribution patterns in the TMJ disc and retrodiscal tissue during mouth opening under different degrees of anterior DDwoR: (A) Relative positional changes among the fossa, disc, and condyle at 0.25s; (B) Equivalent strain distributions and associated morphological alterations at 0.25s; (C) Maximum equivalent strain during mouth opening, and (D) Ratio of the contact area with equivalent strain greater than half-activation strain threshold (ε_0_ = 0.1) to the total surface area of the disc and retrodiscal tissue.

**Fig. 6.**
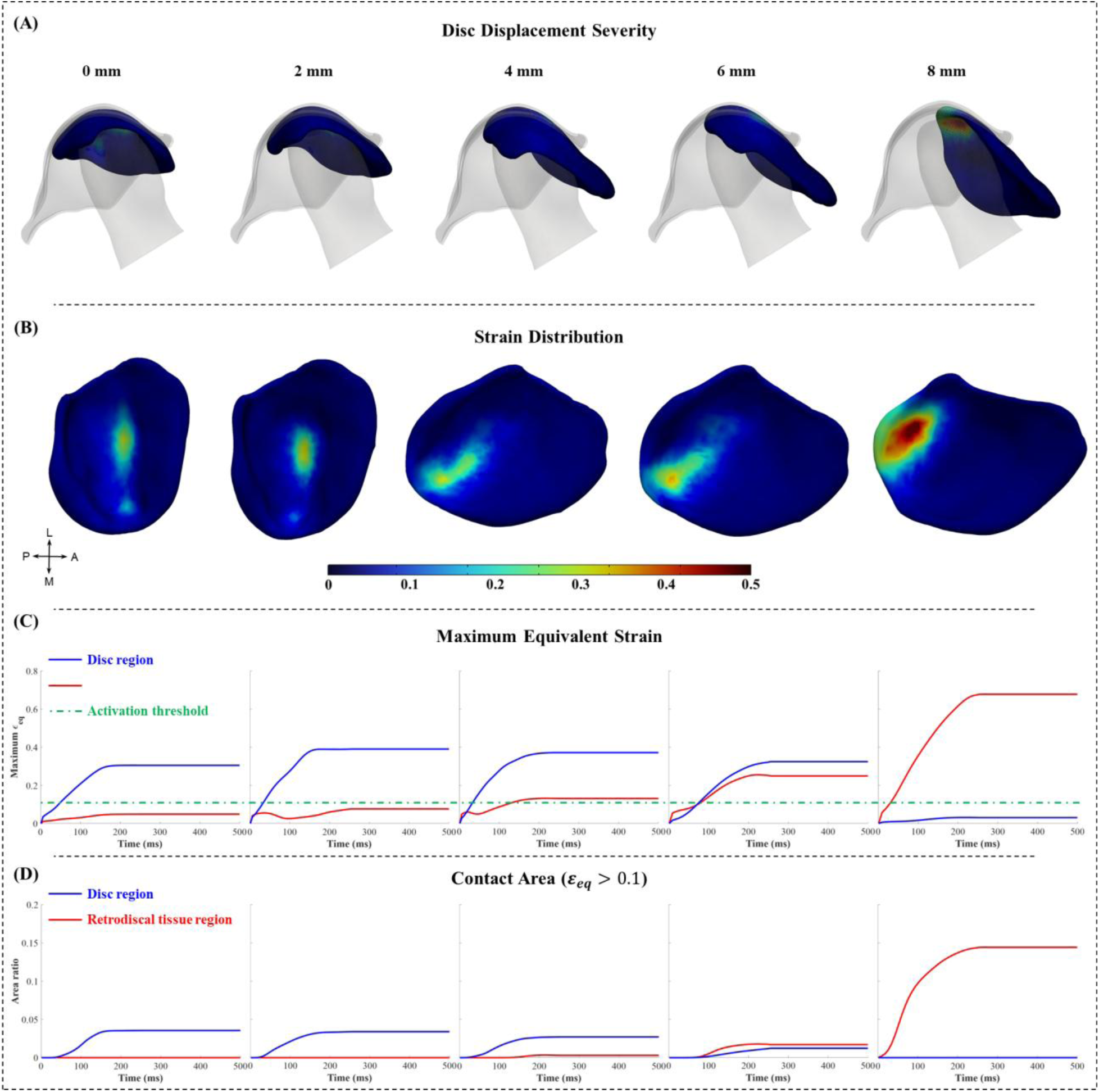
Strain distribution patterns in the TMJ disc and retrodiscal tissue during clenching under different degrees of anterior DDwoR: (A) Relative positional changes among the fossa, disc, and condyle at 0.25s; (B) Equivalent strain distributions and associated morphological alterations at 0.25s; (C) Maximum equivalent strain during mouth opening, and (D) Ratio of the contact area with equivalent strain greater than half-activation strain threshold (ε_0_ = 0.1) to the total surface area of the disc and retrodiscal tissue.

According to the mechanosensitive current formulation in Eqs. 2-3, *ε*_0_ = 0.1 represents the half-activation strain threshold in the Boltzmann-type gating function. Therefore, regions with equivalent strain exceeding 0.1 were considered more likely to activate mechanosensitive channels. **Figs. 5C and 6C** show the maximum equivalent strain in the disc and retrodiscal tissue during maximum mouth opening and clenching. Under the healthy and 2-mm displacement conditions, the maximum equivalent strain in the retrodiscal tissue remained below 0.1, with strain mainly concentrated in the disc. In contrast, when anterior disc displacement exceeded 4 mm, the DDwoR models exhibited maximum equivalent strains greater than 0.1 in the retrodiscal tissue. At 8-mm displacement, the maximum strain occurred in the retrodiscal tissue.

The ratio of the contact area with equivalent strain greater than 0.1 to the total surface area of the disc and retrodiscal tissue is shown in **Figs. 5D and 6D**. When anterior disc displacement exceeded 4 mm, the high-strain contact area progressively increased in the retrodiscal tissue. These findings indicate that displacement greater than 4 mm increased not only the strain magnitude but also the high strain area in the retrodiscal tissue.

**Table 2** summarizes the key strain-related mechanical input parameters, including the maximum equivalent strain in each region, the duration above the half-activation strain threshold of 0.1, and the maximum contact area ratio exceeding this threshold.

**Table 2.** Strain-related mechanical input parameters in the TMJ disc and retrodiscal tissue under different disc displacement conditions.

| Oral task | Parameters | Region | Disc displacement distance (mm) |  |  |  |  |
| --- | --- | --- | --- | --- | --- | --- | --- |
|  |  |  | 0 | 2 | 4 | 6 | 8 |
| Mouth opening | Maximum equivalent strain | Disc | 0.27 | 0.32 | 0.43 | 0.45 | 0.05 |
|  |  | Retrodiscal tissue | 0.05 | 0.04 | 0.11 | 0.40 | 0.65 |
| | Duration above $\varepsilon_0 = 0.1(s)$ | Disc | 0.44 | 0.44 | 0.44 | 0.42 | 0 |
|  |  | Retrodiscal tissue | 0 | 0 | 0.42 | 0.42 | 0.46 |
| | Maximum contact area ratio above $\varepsilon_0 = 0.1(\%)$ | Disc | 2.77 | 1.98 | 2.17 | 2.28 | 0 |
|  |  | Retrodiscal tissue | 0 | 0 | 0.09 | 0.68 | 11 |
| Clenching | Maximum equivalent strain | Disc | 0.30 | 0.38 | 0.37 | 0.32 | 0.03 |
|  |  | Retrodiscal tissue | 0.05 | 0.08 | 0.13 | 0.25 | 0.67 |
| | Duration above $\varepsilon_0 = 0.1(s)$ | Disc | 0.46 | 0.47 | 0.47 | 0.42 | 0 |
|  |  | Retrodiscal tissue | 0 | 0 | 0.38 | 0.42 | 0.48 |
| | Maximum contact area ratio above $\varepsilon_0 = 0.1(\%)$ | Disc | 3.54 | 3.39 | 2.70 | 1.22 | 0 |
|  |  | Retrodiscal tissue | 0 | 0 | 0.28 | 1.77 | 14 |

### 3.2 Action Potentials of Microscopic Nociceptors

In the neural electrophysiological model, the spatial distributions of mechanosensitive ionic current at 0.25 s during maximum mouth opening and clenching are shown in **Figs. 7A and 8A**, respectively. **Figs. 7B and 8B** present the temporal profiles of action potential firing frequency recorded at the soma throughout the simulation, while **Figs. 7C and 8C** show the corresponding membrane current at the soma.

**Fig. 7.**
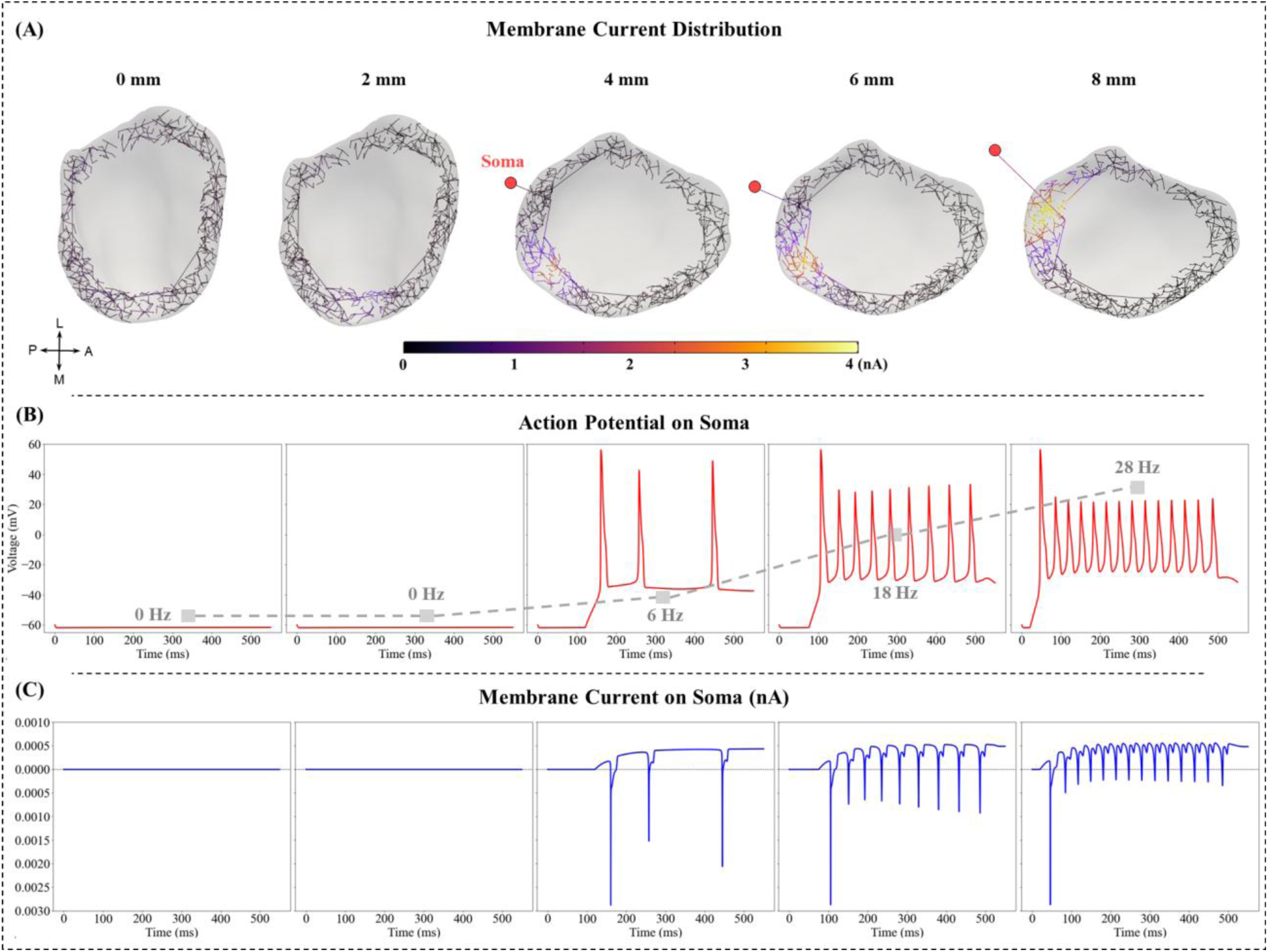
Electrophysiological responses of TMJ nociceptors during mouth opening under different degrees of anterior DDwoR: (A) Spatial distributions of mechanosensitive ionic currents across nociceptor membranes in the TMJ disc and retrodiscal tissue at 0.25 s; (B) Action potential firing frequency recorded at the trigeminal ganglion soma throughout the simulation; and (C) Membrane current responses at the soma throughout the simulation.

**Fig. 8.**
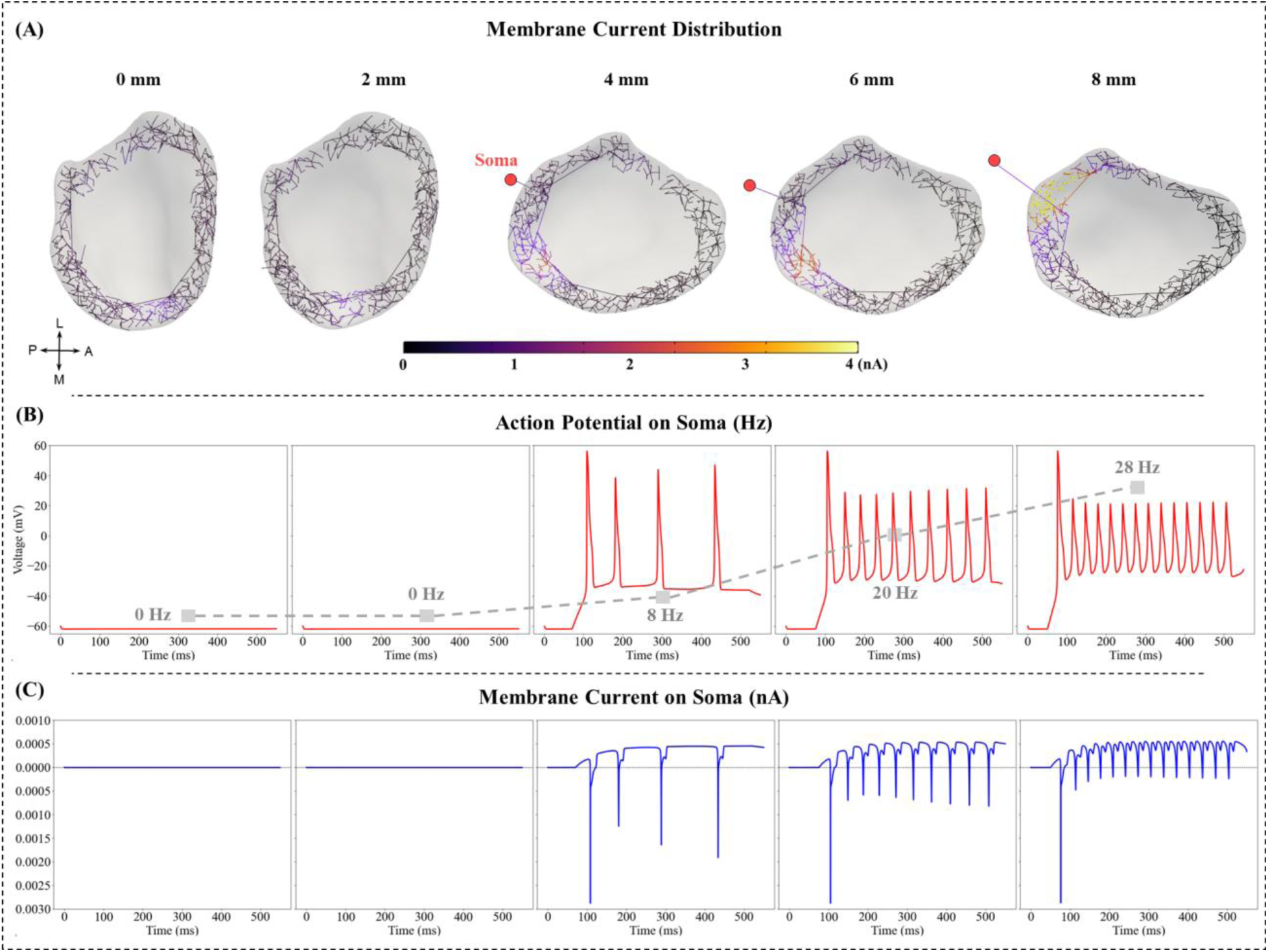
Electrophysiological responses of TMJ nociceptors during clenching under different degrees of anterior DDwoR: (A) Spatial distributions of mechanosensitive ionic currents across nociceptor membranes in the TMJ disc and retrodiscal tissue at 0.25 s; (B) Action potential firing frequency recorded at the trigeminal ganglion soma throughout the simulation; and (C) Membrane current responses at the soma throughout the simulation.

Consistent with the posterior shift and redistribution of the macroscopic strain fields described in **Section 3.1**, increasing anterior DDwoR severity led to larger mechanosensitive current amplitudes and expanded the spatial domain of current generation at nociceptor terminals. Under the healthy and mild displacement conditions, the retrodiscal tissue experienced limited strain activation, resulting in negligible mechanosensitive currents and no action potential firing. In contrast, severe anterior DDwoR (≥4 mm) produced broader terminal depolarization and increased action potential firing at the trigeminal ganglion soma. Firing frequency was used as the primary neural output because it more directly reflects the encoding of nociceptive signal intensity and persistence than voltage amplitude (**Table 3**).

**Table 3.** Action potential firing frequencies under different disc displacement conditions (Hz)

|  | Anterior disc displacement distance |  |  |  |  |
| --- | --- | --- | --- | --- | --- |
|  | 0 mm | 2 mm | 4 mm | 6 mm | 8 mm |
| Mouth opening | 0 | 0 | 6 | 18 | 28 |
| Clenching | 0 | 0 | 8 | 20 | 28 |

## Discussion

This study suggests that anterior DDwoR produces a severity dependent strain pattern characterized by posterior strain redistribution, increased strain magnitude, prolonged activation, and broader retrodiscal tissue involvement, which were associated with enhanced nociceptor activation. These findings support a quantitative structural-mechanical-neural pathway in which altered joint loading may contribute to peripheral nociceptive signaling, suggesting a possible mechanistic link between anterior DDwoR-related changes and pain.

The present results reveal a severity dependent redistribution of strain within the TMJ disc and retrodiscal tissue during mouth opening and clenching. Under the healthy condition, strain was mainly centralized within the disc, suggesting that the disc retained its normal load bearing role between the condyle and the fossa. As anterior displacement increased, the region of elevated strain gradually shifted posteriorly from the disc toward the retrodiscal tissue. This pattern suggests that disc displacement changes the normal load transmission pathway and biomechanical environment in the joint. A key feature of the strain pattern after displacement exceeded 4 mm was the temporal and spatial expansion of strain exceeding the half-activation threshold. The longer duration above the threshold indicates that the retrodiscal tissue was exposed to sustained mechanical input during mandibular function, rather than only transient peak loading. Meanwhile, the increased area ratio shows that strain exceeding this threshold involved a broader region of the retrodiscal tissue. This temporal and spatial expansion indicates that severe DDwoR increases both the persistence and extent of strain activation, which distinguishes larger displacement conditions from healthy and mild displacement conditions and may further increase mechanical input to mechanosensitive nerve endings. Although this assumption has been recognized in previous biomechanical analyses of the TMJ, clinical observations indicate that structural degeneration does not necessarily correspond to pain symptoms; conversely, significant pain can occur in the absence of evident structural changes [4, 11]. A potential explanation for this discrepancy is that nerve distribution and electrophysiological properties are not fully considered in purely biomechanical analyses, although both can modulate neural responses independently of local loading conditions.

To establish a mechanistic link between biomechanics and pain, nociceptors must be explicitly considered. The tissue clearing and 3D nerve mapping techniques [46] provide the anatomical basis for this integration by reconstructing biologically realistic nociceptor networks in the human TMJ disc and retrodiscal tissue. Consistent with previous histological observations, the 3D nerve mapping data confirmed that the central disc is largely devoid of nerves, whereas nerve fibers are concentrated in the peripheral regions, particularly in the anterior and posterior disc regions and retrodiscal tissue. This spatial organization is important because the macroscopic strain results showed that healthy and mild displacement conditions mainly concentrated strain within the disc, whereas larger anterior displacement shifted strain toward the posterior and retrodiscal regions. Thus, the neural consequence of disc displacement depends not only on the magnitude of local strain, but also on whether the strain occurs in sparsely or densely innervated tissue. The electrophysiological model further links these regional strain patterns to nociceptor activation. Mechanosensitive ion channels on nociceptor membranes convert local strain and membrane deformation into depolarizing currents through strain dependent channel gating, modeled here using a Boltzmann type open probability function [61–63]. Compared with previous models that mainly quantified axonal strain or mechanical damage using simplified cylindrical axons [64–66], the present model incorporates ion channel dynamics and therefore directly predicts electrophysiological activity. Firing frequency was emphasized as the main neural output because it better represents nociceptive signal intensity and persistence than voltage amplitude. The gradual decrease in voltage amplitude during repeated firing may reflect changes in membrane excitability during sustained stimulation, including partial inactivation or incomplete recovery of voltage gated sodium channels and the effect of repolarizing potassium currents.

The predicted neural activation under larger displacement conditions can be attributed to the increased spatial overlap between elevated strain and densely innervated retrodiscal tissue. As anterior DDwoR progressed, strain shifted into regions with higher nerve density, thereby increasing mechanosensitive current generation and action potential firing at the trigeminal ganglion soma. This suggests that DDwoR may enhance nociceptive signaling through the coincidence of mechanical loading and peripheral innervation, rather than through structural displacement alone. This interpretation is consistent with previous evidence linking DDwoR pain to compression, stretching, and abnormal loading of the highly innervated and vascularized retrodiscal tissue [25, 67]. Nevertheless, the model output represents peripheral action potential firing at the trigeminal ganglion rather than perceived pain at the brain level, and should therefore be interpreted as an indirect indicator of nociceptive activation potential.

Despite these insights, several limitations should be acknowledged. First, the reconstructed nerve network was derived from porcine innervation data mapped onto human TMJ geometry. While overall anatomical patterns are largely conserved, interspecies differences may affect fine-scale morphological fidelity, highlighting the need for patient-specific nerve mapping approaches in future studies. Second, although the model predicts microscale strains in individual nerve fibers, experimental measurements of cell-scale neural deformation are not currently available for validation, reflecting the difficulty of quantifying in situ cell mechanics. Third, the electrophysiological model employed ion channel parameters obtained from experimentally characterized dorsal root ganglion C-fibers rather than TMJ-specific nociceptors. Given the regional variability in C-fiber channel composition and density, this substitution may introduce inaccuracies in the simulated neural responses. Finally, this study was based on a single individual-specific TMJ model. Using a single controlled TMJ geometry allowed us to isolate the biomechanical and electrophysiological effects of disc displacement severity. Nevertheless, actual DDwoR may involve patient-specific variations in TMJ morphology, occlusal contact, soft tissue properties, and hyoid mobility that were not explicitly modeled. Future studies incorporating multiple subject-specific models, patient-specific imaging data, experimental or in-vivo disc motion validation, and direct validation of TMJ-specific nociceptor activation using in vivo trigeminal afferent recordings or in vitro innervated tissue models are needed to evaluate the robustness and clinical applicability of the proposed framework.

## Conclusion

This study establishes a quantitative multiscale biomechanical-electrophysiological framework linking TMJ structural alterations to peripheral nociceptive activation. By integrating TMJ biomechanics with anatomically informed 3D nociceptor morphology and electrophysiological simulations, the framework quantitatively connected macroscale strain patterns with microscale neural activation. The findings suggest that the spatial overlap between elevated strain and dense retrodiscal innervation may amplify peripheral nociceptive signaling, supporting a potential mechanistic basis for the structure-function-pain relationship in the TMJ under anterior DDwoR conditions.

## CRediT authorship contribution statement

**Jian Chen**: Conceptualization, Data curation, Methodology, Software, Visualization, Writing - original draft, Writing - review and editing. **Shuchun Sun**: Conceptualization, Data curation, Methodology, Writing - review and editing. **Farhad Ahmadi**: Data curation, Methodology, Software, Visualization**. Peng Chen**: Data curation, Methodology, Writing - review and editing. **Jiaxin Chai**: Methodology, Data curation. **Jichao Zhao**: Data curation. **Brooke Damon**: Data curation. **Konstantinia Almpani**: Data curation, Writing - review and editing. **Janice Lee**: Conceptualization, Data curation, Resources, Writing - review and editing. **Hai Yao**: Conceptualization, Methodology, Visualization, Funding acquisition, Project administration, Resources, Supervision, Writing - review and editing.

## Funding sources

This work was funded by the following NIH grants: U01DE031512, R01DE021134, P20GM121342 and R34DE033593.

## Declaration of Competing Interest

The authors declare that they have no known competing financial interests or personal relationships that could have appeared to influence the work reported in this paper.

## Notes

### Competing Interest Statement

The authors have declared no competing interest.

