## Supplementary Materials for "Multiscale Biomechanical and Electrophysiological Modeling of Nociceptive Activation in Musculoskeletal Joint Disorders: Insights from the Temporomandibular Joint"

**Supplementary Note 1: Hill-Type Muscle Model Formulation and Parameters**

The masticatory muscles were modeled as point-to-point Hill-type actuators. The total force $F^{total}$ generated by each muscle is calculated as the sum of active contractile force and passive elastic force. Our implementation follows the Hill-type formulation tailored for jaw biomechanics [1-3] :

$$F^{total} = F^{max} \cdot [a \cdot f_{active}(\tilde{l}) + P_{frac} \cdot f_{passive}(l)] \tag{1}$$

where $F^{max}$ is the maximum muscle force (N), $a$ is the muscle activation level ($0 \leq a \leq 1$), $f_{active}$ is the active force component, $\tilde{l}$ is the normalized fiber length, $P_{frac}$ is the passive fraction coefficient. This value is set as 0.015 [1], $f_{passive}$ is the passive elastic component, $l$ is the total muscle length, including the tendon.

Maximum Force $F^{max}$ is calculated as the product of the muscle's Physiological Cross-Sectional Area ($PCSA$) and a specific tension constant of 40 N/cm² [2] .

The active force component $f_{active}$ describes the force generation capacity of the sarcomeres as a function of normalized fiber length $\tilde{l}$. The model utilizes a cosine-based approximation of the force-length curve [1]:

$$f_{active}(\tilde{l}) = \begin{cases} 0.5 \cdot (1 + \cos(2\pi \cdot \tilde{l})) & if\ 0.5 < \tilde{l} < 1.5 \\ 0 & otherwise \end{cases} \tag{2}$$

The normalized fiber length $\tilde{l}$ is calculated by subtracting the tendon length $L^{tendon}$ from the current total muscle length $l$ and normalizing by the optimal fiber length $L^{opt}_{fiber}$ [1]:

$$\tilde{l} = \frac{l - L^{tendon}}{L^{opt}_{fiber}},\ L^{tendon} = L^{opt} \cdot R_{tendon} \tag{3}$$

where $L^{opt}$ is the total muscle length at which active force is maximal, and $R_{tendon}$ is the tendon-to-muscle length ratio.

The passive elastic component $f_{passive}$ represents the resistance of the muscle tissue to stretching. In our implementation, a linear ramp function was used to approximate the passive curve [1]:

$$f_{passive}(l) = \begin{cases} 0 & if\ l \leq L^{opt} \\ \frac{l - L^{opt}}{L^{max} - L^{opt}} & if\ L^{opt} < l < L^{max} \\ 1.0 & if\ l \geq L^{max} \end{cases} \quad (3)$$

where $L^{max}$ is the maximum muscle length. The parameters governing the muscle model were assigned according to **Table 1** [3].

**Table 1:** Literature-derived parameters of the Hill-type muscle actuators that are constant across subjects

| Muscle Name | $PCSA\ (cm^2)$ | $F^{max}(N)$ | $L^{opt}(mm)$ | $L^{max}(mm)$ | $R_{tendon}$ |
|---|---|---|---|---|---|
| Anterior Temporalis | 3.95[2] | 158.0[2] | 75.54[2] | 95.92[2] | 0.50[2] |
| Middle Temporalis | 2.39[2] | 95.6[2] | 65.81[2] | 93.36[2] | 0.48[2] |
| Posterior Temporalis | 1.89[2] | 75.6[2] | 77.11[2] | 101.08[2] | 0.51[2] |
| Superficial Masseter | 4.76[2] | 190.4[2] | 51.46[2] | 66.88[2] | 0.46[2] |
| Deep Masseter | 2.04[2] | 81.6[2] | 29.07[2] | 44.85[2] | 0.29[2] |
| Medial Pterygoid | 4.37[2] | 174.8[2] | 40.51[2] | 50.63[2] | 0.64[2] |
| Inferior Lateral Pterygoid | 1.67[2] | 66.9[2] | 31.5[1] | 41.5[1] | 0.00[a][1] |
| Superior Lateral Pterygoid | 0.72[b][1] | 28.7[b][1] | 27.7[1] | 37.7[1] | 0.00[a][1] |
| Anterior Digastric | 1.00[2] | 40.0[2] | 31.24[c] | 45.10[c][1] | 0.00[a][1] |
| Anterior Mylohyoid | 0.89[d][1] | 35.4[1] | 25.08[c] | 45.10[c][1] | 0.00[a][1] |
| Posterior Mylohyoid | 0.89[d][1] | 35.4[1] | 27.13[c] | 45.10[c][1] | 0.00[a][1] |
| Geniohyoid | 0.80[3] | 32.0[1] | 31.79[c] | 45.10[c][1] | 0.00[a][1] |

[a] In our implementation, $R_{tendon}$ was assigned to be 0 for these muscles [1].

[b] Modeled as 30% of the total Lateral Pterygoid capacity, with the Inferior head representing the other 70%, derived from Peck et al. [2].

[c] In this implementation, the $L^{opt}$ was adjusted to achieve the maximum possible mouth opening and $L^{max}$ was assigned to be the same for the Anterior Digastric, Anterior and Posterior Mylohyoid, and Geniohyoid muscles.

[d] The total Mylohyoid ($PCSA$ = 1.77 $cm^2$ [4]) was divided equally between the Anterior and Posterior segments.

The TMJ disc and retrodiscal tissue were assigned a density of $1.05\ \mathrm{g/cm^3}$, adopted from the experimentally measured wet tissue density of cartilaginous lumbar annulus fibrosus reported by Yao et al [5]. In that study, the annulus fibrosus had a water content above 80%, indicating that the reported density represents a highly hydrated cartilaginous tissue and is therefore applicable as an approximate density for the TMJ disc, which is also a water-rich fibrocartilaginous tissue. Kuo et al. reported an average tissue water volume fraction of approximately 80% in the human TMJ disc [6], further supporting this assumption. The same density was assigned to the retrodiscal tissue.

The mandible was assigned a mass of $200\ \mathrm{g}$, consistent with the reported dynamic jaw model [1]. The skull and hyoid bone were modeled as fixed rigid bodies; therefore, their masses did not contribute to the mouth opening or clenching simulations. The articular cartilages, disc attachments, anatomical ligaments, and muscle actuators were not assigned separate density values because they were represented using non-volumetric formulations rather than volumetric finite element bodies.

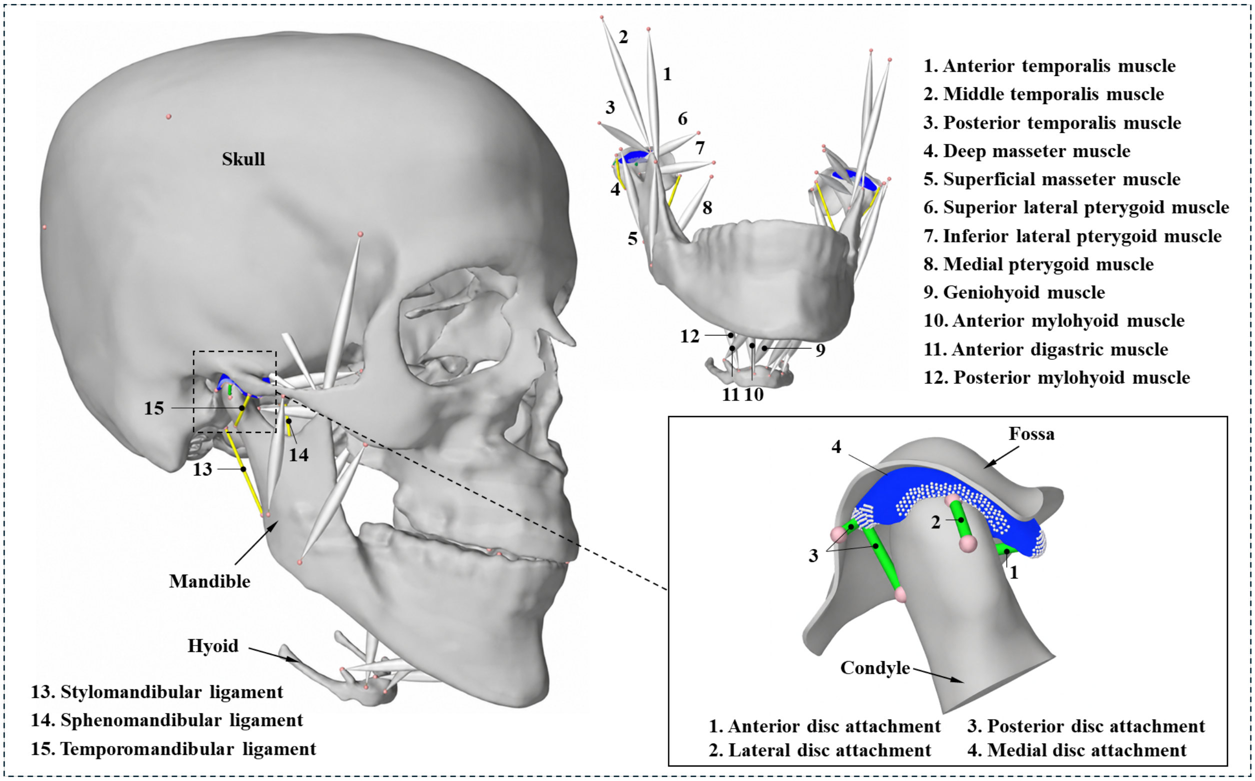


**Supplementary Figure 1:** Musculoskeletal finite element model of the human jaw. The model includes the skull, mandible, hyoid bone, twelve jaw muscles, and three anatomical TMJ ligaments: the stylomandibular, sphenomandibular, and temporomandibular ligaments. Four disc attachment constraint groups were included in the anterior, posterior, lateral, and medial directions. The medial attachment is located opposite the lateral attachment and is therefore not visible in this view.

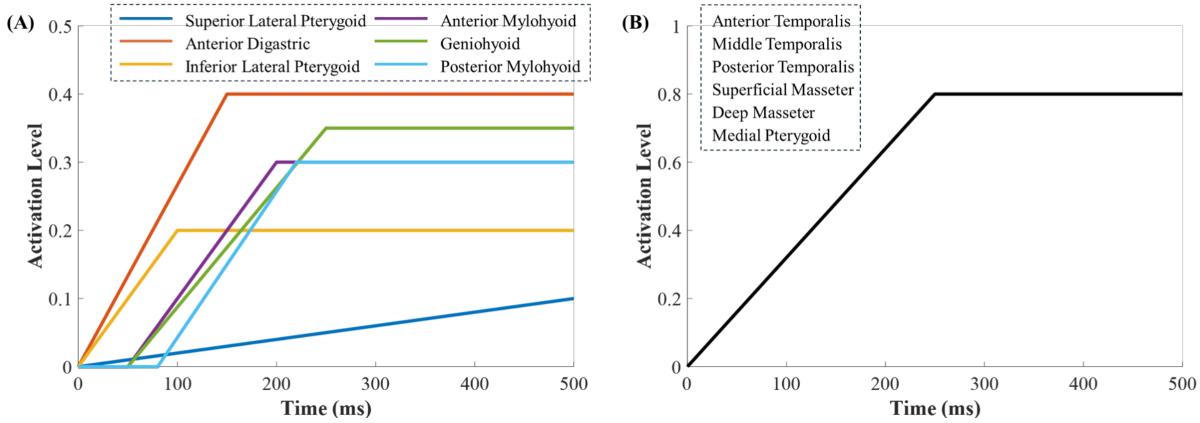


**Supplementary Figure 2:** Muscle activation levels for TMJ biomechanical model: (A) Muscle activation levels during mouth opening. (B) Muscle activation levels during clenching. For the mouth opening simulation, activation levels were selected to reproduce the target maximum mouth opening behavior. For the clenching simulation, muscle activation levels were selected to generate a physiologically reasonable TMJ joint reaction force of approximately 40 N [7] These task-based prescriptions follow the modeling strategy used in previous dynamic jaw simulations, in which muscle activations were computationally determined according to the simulated oral task [1].

**Supplementary Materials Note 2: Kinematic data collection and model validation**

For tracking skull and mandible movement, four high-speed infrared cameras (Prime 13, Optitrack, OR) mounted on a tripod were equipped. The tripod was positioned approximately 1m from the subject, providing sufficient space for examiner movement while maintaining high tracking accuracy. The cameras were arranged with a non-collinear configuration to reduce capture errors. Prior to data collection, the cameras were calibrated with a calibration wand to determine their positions and orientations in a common coordinate system. After calibration, the camera data were used to reconstruct the 3D trajectories of 5mm reflective markers mounted on rigid frames. To ensure accurate rigid-body motion capture, seven reflective markers were assembled on a skull frame. The frame was tightly fixed to the subject's head during both motion capture and CBCT imaging, thereby enabling registration between the motion capture system and CBCT coordinates. Six reflective markers were assembled on a mandible frame. This frame was connected to the teeth via a disposable bracket attached to the buccal surface of the anterior mandibular arch using a two-part bis-acrylic polymer (Tempsmart, GC America). Detailed photographs of the setup are provided in the **Supplementary Figure 3**.

Kinematic analysis was performed for maximum mouth opening. During the 0.25s transition from the closed mouth position to maximum mouth opening, simulated mandibular motion was compared with experimentally acquired kinematic data under healthy conditions without disc displacement. The validation included assessments of mandibular position and configuration, central incisor displacement, and rotation magnitude, as shown in **Supplementary Figure 4**. Central incisor displacement and mandibular rotation magnitude were compared over the 0.25 s mouth opening interval. These two metrics were selected because they represent the primary translational and rotational components of mandibular opening and are directly relevant to the boundary motion driving the TMJ biomechanical simulation. For central incisor displacement, the simulated endpoint closely matched the experimental endpoint, with an endpoint absolute error of 0.25 mm and a relative endpoint error of 0.76%. The Pearson correlation coefficient was 0.96, indicating a highly similar temporal trend between simulation and experiment. The normalized root mean square error was 14.80%, suggesting moderate time dependent discrepancies during the opening process. For mandibular rotation magnitude, the endpoint absolute error was 2.00°, corresponding to a relative endpoint error of 10.77%. The Pearson correlation coefficient was 0.94, indicating strong temporal agreement, while the normalized root mean square error was 15.26%, also reflecting moderate time dependent discrepancies. Overall, the model reproduced the endpoint mandibular opening configuration and the general temporal trends of mouth opening, although deviations remained during the intermediate phase of motion.

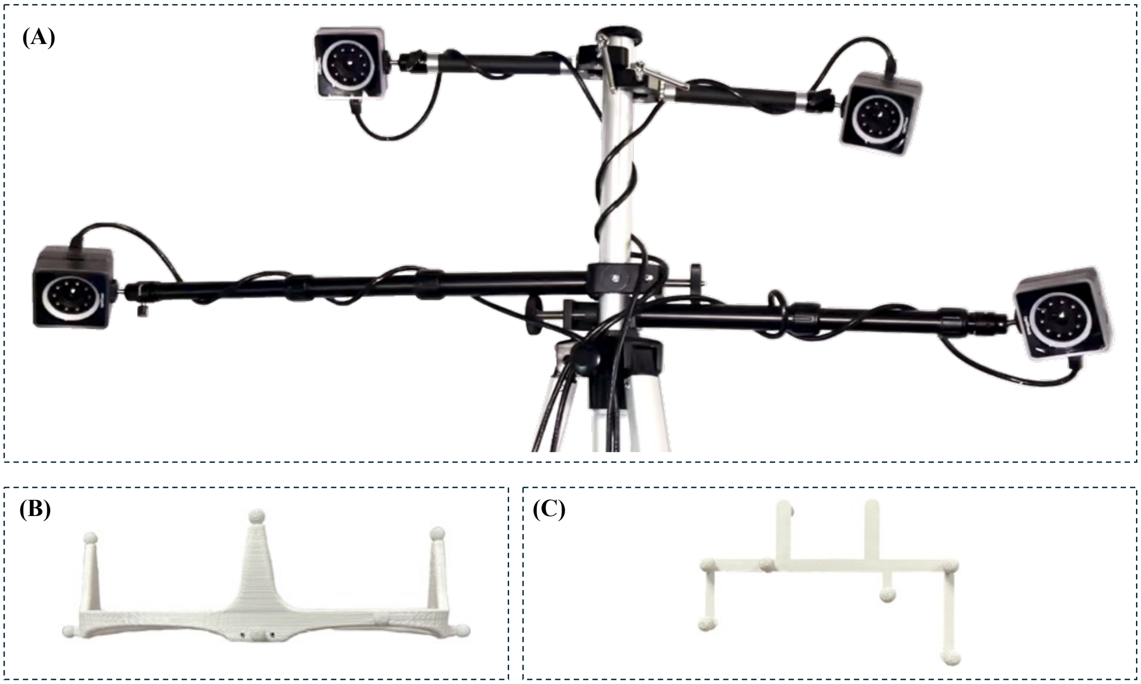


**Supplementary Figure 3.** Motion capture setup for tracking skull and mandible movement. (A) High-speed cameras mounted on tripods. (B) Skull marker frame with reflective markers. (C) Mandible marker frame with reflective markers.

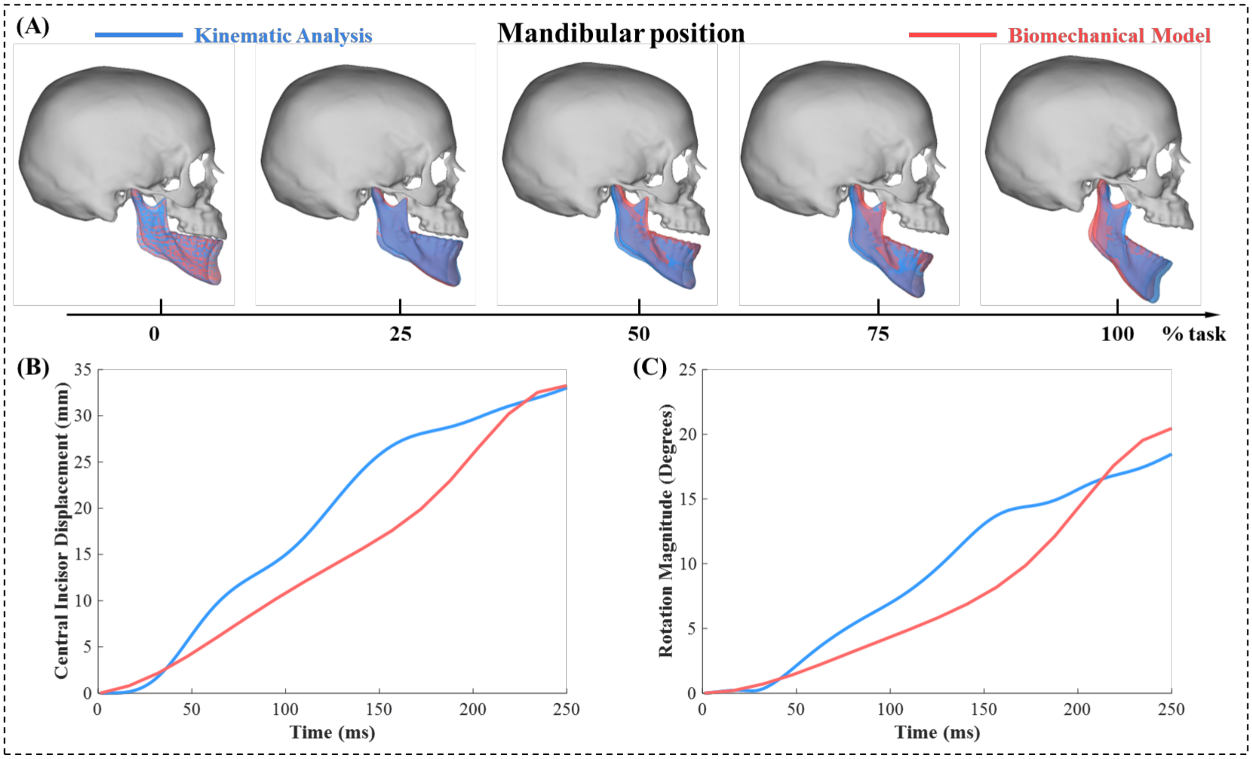


**Supplementary Figure 4.** Experimental-simulated comparison of mandibular kinematics under healthy conditions. (A) Mandibular position and configuration. (B) Central incisor displacement. (C) Rotation magnitude.

In addition, simulated mandibular motion was compared across varying degrees of disc displacement to evaluate whether disc displacement severity altered mandibular kinematics, as shown in **Supplementary Figure 5**. Mandibular motion remained nearly unchanged across different severities of disc displacement.

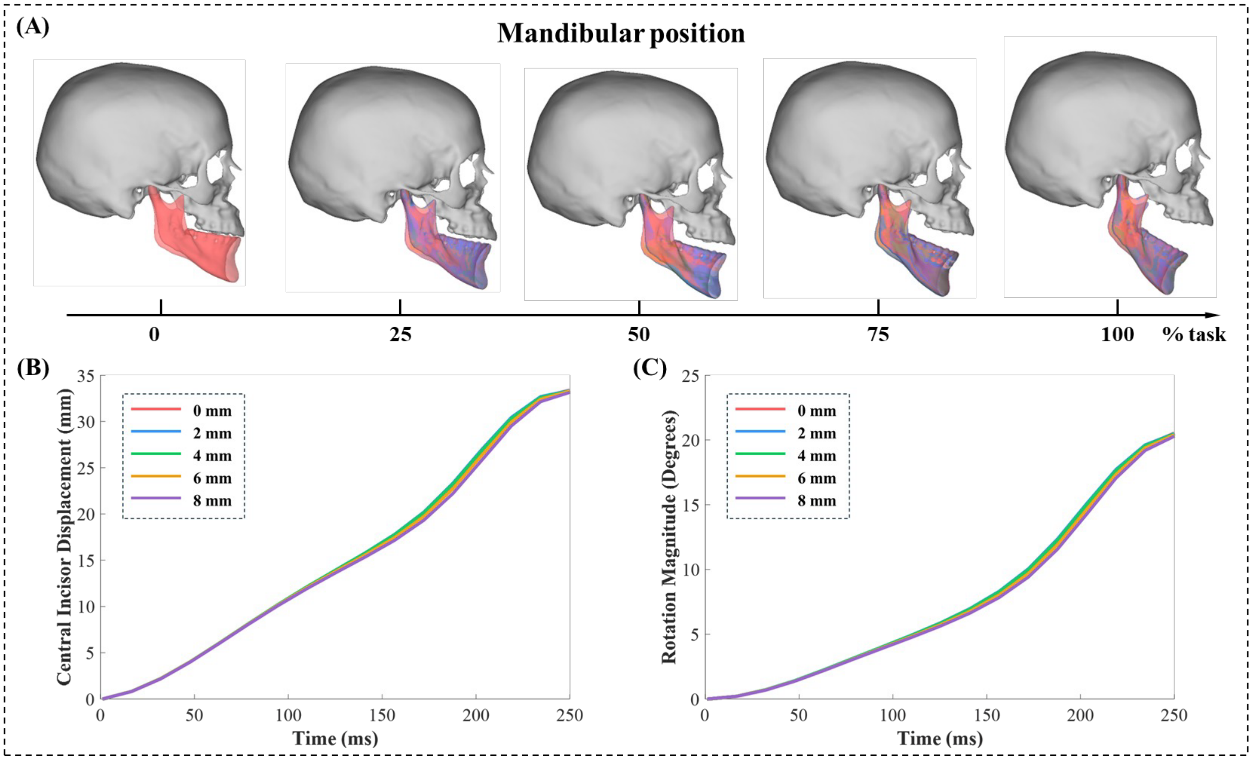


**Supplementary Figure 5.** Comparison of mandibular kinematics across simulated disc displacement conditions. (A) Mandibular position and configuration. (B) Central incisor displacement. (C) Rotation magnitude.

**Supplementary Materials Note 3: Neural Electrophysiological Model**

The membrane model of the nociceptor contains Hodgkin–Huxley-type ion channels, which is represented as follows:

$$C_m \frac{dV}{dt} + \sum_{\alpha=ion} I_\alpha + I_{MS} = 0 \quad (4)$$

where $V$ is the membrane potential of the nociceptors, $C_m$ is the membrane capacitance per unit area, $I_{MS}$ is the ionic current of the strain-activated mechanosensitive ion channels, served as the driving input for neural activation in our membrane model, $I_\alpha$ stand for the ionic current driven by the potential difference and governed by the conductance, shown as follows:

*Tetrodotoxin (TTX)-sensitive sodium channels:*

$$I_{Nattxs} = \bar{g}_{Nattxs} m^3 h (V - E_{Na}) \quad (5)$$

$$\tau_m(V)\frac{dm}{dt} = m_\infty(V) - m \qquad \tau_h(V)\frac{dh}{dt} = h_\infty(V) - h$$

$$m_\infty(V) = \frac{\alpha_m(V)}{\alpha_m(V) + \beta_m(V)} \qquad h_\infty(V) = \frac{\alpha_h(V)}{\alpha_h(V) + \beta_h(V)}$$

$$\tau_m(V) = \frac{1}{\alpha_m(V) + \beta_m(V)} \qquad \tau_h(V) = \frac{1}{\alpha_h(V) + \beta_h(V)}$$

$$\alpha_m(V) = \frac{17.235}{1 + \exp\left(\frac{V + 7.58}{-11.47}\right)} \qquad \alpha_h(V) = 0.23688 \exp\left(-\frac{V + 115}{46.33}\right)$$

$$\beta_m(V) = \frac{17.235}{1 + \exp\left(\frac{V + 66.2}{19.8}\right)} \qquad \beta_h(V) = 10.8 \exp\left(\frac{V - 11.8}{-11.998}\right)$$

where $\bar{g}_{Nattxs} = 0.001699\, S/cm^2$ is maximal ionic conductance, $m$ and $h$ represent gating variables and the rest gating variables follow the same formulation, $E_{Na} = 60\, mV$ is the reversal potential for sodium ions.

*TTX-sensitive persistent sodium channels:*

$$I_{NaP} = \bar{g}_{NaP} m^3 (V - E_{Na}) \tag{6}$$

$$\alpha_m(V) = \frac{17.235}{1 + \exp\left(\frac{V + 27.58}{-11.47}\right)} \qquad \beta_m(V) = \frac{17.235}{1 + \exp\left(\frac{V + 86.2}{19.8}\right)}$$

where $\bar{g}_{NaP} = 0.00005\ S/cm^2$, $E_{Na} = 60\ mV$.

*Nav1.9 TTX- resistant sodium channels:*

$$I_{NaV1.9} = \bar{g}_{NaV1.9} mh (V - E_{Na}) \tag{7}$$

$$\alpha_m(V) = \frac{1.548}{1 + \exp\left(\frac{V - 11.01}{-14.871}\right)} \qquad \alpha_h(V) = \frac{0.2574}{1 + \exp\left(\frac{V + 63.264}{3.7193}\right)}$$

$$\beta_m(V) = \frac{8.685}{1 + \exp\left(\frac{V + 112.4}{22.9}\right)} \qquad \beta_h(V) = \frac{0.53984}{1 + \exp\left(\frac{V + 0.27853}{-9.0933}\right)}$$

where $\bar{g}_{NaV1.9} = 0.00005\ S/cm^2$, $E_{Na} = 60\ mV$.

*Nav1.8 TTX- resistant sodium channels:*

$$I_{NaV1.8} = \bar{g}_{NaV1.8} m^3 h (V - E_{Na}) \tag{8}$$

$$\alpha_m(V) = \frac{3.83}{1 + \exp\left(\frac{V + 2.58}{-11.47}\right)} \qquad \alpha_h(V) = 0.013536 \exp\left(-\frac{V + 105}{46.33}\right)$$

$$\beta_m(V) = \frac{6.894}{1 + \exp\left(\frac{V + 61.2}{19.8}\right)} \qquad \beta_h(V) = \frac{0.61714}{1 + \exp\left(\frac{V - 21.8}{-11.998}\right)}$$

where $\bar{g}_{NaV1.8} = 0.02\ S/cm^2$, $E_{Na} = 60\ mV$.

*Delayed rectifier potassium channels:*

$$I_{Kdr} = \bar{g}_{Kdr} n(V - E_K) \tag{9}$$

$$\alpha_n(V) = \frac{0.001265(V + 14.273)}{1 - \exp\left(\frac{V + 14.273}{-11.47}\right)} \qquad \beta_n(V) = 0.125\exp\left(\frac{V + 55}{-2.5}\right)$$

where $\bar{g}_{Kdr} = 0.000878\ S/cm^2$, $E_K = -85\ mV$.

*A-type potassium channels:*

$$I_{KA} = \bar{g}_{KA} m^4 h(V - E_K) \tag{10}$$

$$\alpha_m(V) = \frac{1.4}{1 + \exp\left(\frac{V + 7}{-12}\right)} \qquad \alpha_h(V) = \frac{0.0175}{1 + \exp\left(\frac{V - 30}{8}\right)}$$

$$\beta_m(V) = \frac{0.49}{1 + \exp\left(\frac{V + 10}{4}\right)} \qquad \beta_h(V) = \frac{1.3}{1 + \exp\left(\frac{V + 7}{-10}\right)}$$

where $\bar{g}_{KA} = 0.0012\ S/cm^2$, $E_K = -85\ mV$.

*Kv7/M channels:*

$$I_{KM} = \bar{g}_{KM} m(V - E_K) \tag{11}$$

$$m_\infty(V) = A_2 + (A_1 - A_2)\frac{1}{1 + \exp\left(\frac{V + 42}{12}\right)} \qquad \tau_m(V) = 60 + \frac{\beta_m(V)}{0.009(1 + \alpha_m(V))}$$

$$\alpha_m(V) = \exp\left[0.2646(V + 42)\right] \qquad \beta_m(V) = \exp\left[0.10584(V + 42)\right]$$

where $\bar{g}_{KM} = 0.0001\ S/cm^2$, $E_K = -85\ mV$, $A_1 = -0.07245$, $A_2 = 1.13462$.

*Hyperpolarization-activated channels:*

$$I_h = \bar{g}_h l(V - E_h) \tag{12}$$

$$l_\infty(V) = \frac{1}{1 + \exp\left(\frac{V + 73}{10}\right)} \qquad \tau_l(V) = \frac{\beta_l(V)}{0.02(1 + \alpha_l(V))}$$

$$\alpha_l(V) = \exp\left[0.08116(V + 75)\right] \qquad \beta_l(V) = \exp\left[0.033264(V + 75)\right]$$

where $\bar{g}_h = 0.0001\ S/cm^2$, $E_h = -40\ mV$.
